# Sampling bias and uneven data coverage shape observed biodiversity patterns in a megadiverse island archipelago hotspot

**DOI:** 10.1101/2025.09.13.676052

**Authors:** Kier Mitchel E. Pitogo, Camila G. Meneses, Syrus Cesar P. Decena, Christian E. Supsup, Hannah E. Som, Justin M. Bernstein, Kin Onn Chan, Mark William Herr, Rafe M. Brown

**Author notes:** Corresponding author: Kier Mitchel E. Pitogo. Department of Integrative Biology & MSU Museum, Michigan State University, East Lansing, MI, USA.

## Abstract

Where and how species are sampled can shape biodiversity knowledge, spatial patterns, and data-driven conservation. In megadiverse regions of the Global South, sampling remains uneven, and available data often lack synthesis needed to assess spatial heterogeneity in recorded biodiversity for conservation planning. This limitation is also evident in areas designated for biodiversity conservation. We examine this in the Philippines, where long-term research effort and policy development have supported expansion of conservation-relevant areas (hereafter “Philippine conservation-relevant areas” or PCAs). Using over a century of digitally accessible species occurrence records compiled and manually curated, we analyzed the country-wide spatial distribution of observed amphibian and squamate reptile species richness. We further tested whether observed richness across PCAs is associated with area, topographic relief, and sampling effort, and whether these relationships differ between protected and conserved areas (with management or governance schemes) and key biodiversity areas (unmanaged but priority areas for conservation). Results show strong spatial heterogeneity in occurrence records, with preserved specimens comprising 92% of all records and strongly influencing observed richness patterns. Citizen-science data tend to reinforce already well-sampled regions, whereas literature-derived records provide additional coverage in comparatively undersampled areas. PCAs show variable representation in the dataset, with observed richness and sampling effort generally increasing with area. However, this richness–area relationship weakens with increasing topographic relief. Differences in predictor–richness relationships between PCA types are also evident, with stronger area and sampling effects in protected and conserved areas and a relatively stronger elevational effect in key biodiversity areas. Notably, several high-richness areas occur outside formally designated protected areas. These results highlight uneven biodiversity data coverage and suggest that observed richness patterns may reflect both sampling structure and spatial variation. We emphasize integrating multiple biodiversity data sources and identifying undersampled regions to improve future biodiversity assessments.

## Introduction

Biodiversity occurrence data remain unevenly distributed across taxonomic groups and geographic regions, with pronounced biases against global biodiversity hotspots [1–5]. Several factors contribute to these biases, including limited research capacity [6,7], preferential focus on certain taxonomic groups [3,4,8] and charismatic taxa [9,10], ease of site accessibility [11–13], and prevailing tendency to conduct research in areas of special interest [14,15]. As a result, many regions and their wildlife remain poorly sampled—and where data exist, they are often not digitally accessible or fall into “biodiversity blind spots”, shaping biodiversity patterns [5]. These sampling biases can hinder robust macroecological analyses and obstruct evidence-based conservation planning [17–20], particularly at national scales where most conservation policies and management actions are implemented [21–23]. Therefore, identifying where biodiversity data exist, for which taxa, and to what extent is essential for proactively addressing these biases and strengthening the knowledge base needed for effective conservation.

Challenges in biodiversity data availability are persistent even within global biodiversity hotspots and megadiverse countries, like the Philippines, where gaps and biases in data coverage may skew biodiversity knowledge toward certain regions and taxonomic groups [24–26]. As a megadiverse and biogeographically unique country threatened by habitat loss [27,28], the Philippines urgently needs robust and data-informed conservation strategies. However, limited availability and inequitable discoverability of biodiversity data, and a lack of information on how these data are spatially distributed, constrains efforts to evaluate sampling completeness and to guide national biodiversity strategies and research priorities [23,29]. The need for synthesized biodiversity data to inform conservation planning is especially urgent, given ongoing biodiversity loss and the global momentum under the Kunming–Montreal Global Biodiversity Framework [30,31]. These factors have prompted policy responses to expand protected areas and recognize other effective area-based conservation measures, which now cover approximately 15% of the Philippines’ land area [32]. This expansion not only demands effective management grounded in ecologically representative, well-connected, and equitably governed networks of protected and conserved areas [30] but also must deliver measurable, positive outcomes for biodiversity [33]. A critical first step towards achieving such outcomes is to strengthen biodiversity data by improving sampling coverage and accessibility [21,23,29,34,35–37].

A key step toward ensuring that biodiversity knowledge is sufficient to support conservation strategies is to assess where biodiversity data are spatially distributed and how these patterns align with areas considered important for biodiversity (hereafter ‘Philippine conservation-relevant areas’ or PCAs), including protected and conserved areas (with governance and management schemes) and key biodiversity areas (priority areas for conservation, with no management or governance schemes) [38]. Such an approach remains lacking for the Philippines despite long-standing efforts to document wildlife in this country [24–26,39–48]. Although data deficiencies affect many taxonomic groups, some are better represented due to sustained research and collection efforts. Amphibians and squamate reptiles (herpetofauna) offer a compelling case for such analysis in the Philippines. As two of the most consistently documented vertebrate taxa in the country, herpetofauna provide a uniquely comprehensive dataset for assessing spatial documentation patterns. While acknowledging their distinct ecological niches, we integrate these groups as a singular, informative lens to evaluate biodiversity records across varying sampling landscapes. This group has benefited from over a century of active and sustained research effort [25,39]; in fact, recent years have seen an acceleration of engagement in amphibian and reptile studies—marked by an increasing number of researchers, broader investigation scopes, and expanding publication output [25]. Such growth underscores the timeliness of using herpetofauna to model biodiversity data coverage. Furthermore, endemic herpetofauna are relatively well-sampled genetically, with voucher specimens housed in museum collections [26] and curated occurrence records are available in substantial volume across peer-reviewed literature [49,50]. Because herpetofauna serve as effective surrogates for vertebrate biodiversity patterns in KBAs across the country [51], they provide a robust framework to evaluate spatial patterns in biodiversity data, ensuring that country-wide conservation efforts are grounded in the best available evidence.

Leveraging species occurrence data on Philippine herpetofauna spanning ∼125 years (1900s–2025) that are made digitally accessible, we compiled, manually curated, and analyzed occurrence records to conduct country-wide spatial assessments of observed species richness of amphibians and squamate reptiles using data from multiple sources. Specifically, we ask: (1) How is observed herpetofaunal species richness spatially distributed across the Philippines? (2) How do these patterns differ across datasets derived from museum collections, citizen-science platforms, and peer-reviewed literature? (3) How does area, topographic relief, and sampling intensity shape variations in observed species richness across PCAs? And (4) do these variations in observed species richnness differ between protected and conserved areas and key biodiversity areas? Our findings offer long-overdue yet novel insights into spatial distribution of biodiversity data in the Philippines, particularly in relation to PCAs. Drawing from our results, we propose and discuss enabling mechanisms to improve primary biodiversity data collection, strengthening the knowledge base necessary in assessing management effectiveness for area-based conservation measures needed in the country [52,53]. By identifying key data shortfalls and sampling biases, this study contributes to the growing body of empirical evidence underscoring the critical role of integrating digitally accessible biodiversity data from multiple sources in revealing sampling gaps and biases, guiding future studies towards poorly sampled areas and informing conservation priorities.

## Materials & methods

### Data assembly and curation

All georeferenced occurrence records for amphibians and squamate reptiles from the Philippines were downloaded from the Global Biodiversity Information Facility (GBIF) on 31 January 2025 (amphibians) and 07 February 2025 (squamate reptiles). To reduce data duplication, we initially excluded records categorized as material citations and retained only human-observation records from iNaturalist (research grade) and HerpWatch Pilipinas, a non-government organization from the Philippines that documents herpetofaunal diversity. These data were then merged with expert-curated species occurrence data from the literature: amphibians from the Amphibians of the Philippines, part I: checklist of the species [49] and snakes from the Synopsis of snakes of the Philippines [50]. To avoid overlap between GBIF records and these curated datasets (also sourced from GBIF), only GBIF records dated post-publication—2015 onwards for amphibians and 2018 onwards for snakes—were retained, since GBIF records for these groups collected before 2015 (for amphibains) and 2018 (for snakes) had already undergone rigorous curation. For lizards, all GBIF records were included since no curated datasets are available. Duplicate specimen records across institutions were identified using shared field numbers. When duplicates were detected, only records from the University of Kansas Natural History Museum (KU) were retained. Finally, we incorporated verifiable occurrence records of non-museum-catalogued specimens from peer-reviewed literature compiled by a recent review on Philippine herpetology over the last 20 years [25], along with a few additional entries. For consistency in categorization, all museum records were treated as “preserved specimen” records; iNaturalist and HerpWatch Pilipinas as “citizen science” records; and data from peer-reviewed literature as “material citation” records.

After initial clean-up, we recovered 471 nominal and candidate species (i.e., divergent populations supported by evidence from peer-reviewed studies but not yet formally described) of terrestrial amphibians and squamate reptiles that have digitally accessible occurrence records. Occurrence records for each species were then manually curated using currently accepted taxonomic treatments and synonyms, following the AmphibiaWeb [54] for amphibians and The Reptile Database [55] for reptiles. Each species’ occurrence records were mapped in QGIS 3.4 Madeira [56] to assess spatial accuracy of each point. Records falling outside the known geographic range of a species were excluded, following genetically confirmed species boundaries from published literature summarized in the recent Philippine herpetology review paper [25], and those lacking verifiable documentation (e.g., curated vouchered specimens, occurrences within known distribution from peer-reviewed literature). In cases where a formerly widespread species had been taxonomically split, historical records were reassigned to currently accepted species name based on updated diagnostic and distributional information in taxonomic studies. For species with unclear taxonomic or geographic boundaries, only records supported by verifiable documentation were retained. Since expert-curated distributional literature for Philippine lizards is lacking, we adopted an additional measure by comparing our preliminary range estimates with the Global Assessment of Reptile Distributions [57–58]. This conservative step was taken to minimize potential overestimation of species richness.

Records not identified to the species level were excluded unless a species name or a placeholder name (e.g., sp. + island name, sp. + placeholder number) could be confidently assigned without the risk of double-counting (e.g., when only a single species from a genus is known to occur in the area). For example, the frog genus *Platymantis* remains taxonomically unresolved with many potentially undescribed species [59–61]. We accounted for this uncertainty by adopting a conservative approach by including records of candidate species occurring in areas with no known range overlap with closely related congeners. This strategy minimizes the risk of double-counting should these populations ultimately represent a single species, while also retaining valuable data that would otherwise be discarded. This decision was informed by phylogenetic evidence, following published phylogeny of Ceratobatrachidae [59].

All records lacking locality information were discarded. Additionally, records with coordinates at 16.45°N, 120.55°E and 13°N, 122°E from GBIF were excluded, as the associated locality descriptions did not match the projected geographic locations. These coordinates appear to be generalized placeholders rather than accurate site data. In the final dataset, we retained records whose geographic coordinates were consistent with their general (provincial-level) locality. Although minor spatial imprecision is unlikely to affect broad, country-scale patterns, it may influence species richness estimates within individual PCAs used in the modeling. However, any resulting bias is expected to be minimal because most PCAs are supported by numerous occurrence records, species are represented by multiple points within PCAs due to the integration of multiple data sources, and the overall sample size is large (n = 316 PCAs).

## Statistical analyses

### Mapping herpetofaunal diversity across the Philippines

To assess the spatial distribution of observed herpetofaunal species richness patterns, we aggregated occurrence records within a spatial grid and mapped resulting species richness values. Occurrence records for amphibians and squamate reptiles were used to generate a presence–absence matrix (PAM) at a 10-km grid resolution across the Philippines, using prepare_base_pam function from the R package biosurvey [62]. Based on the overall geographic template of the Philippines, we selected a 10-km grid resolution for this visual assessment because it better captures coarse-scale patterns across islands of varying sizes. Smaller grids (e.g., 5 km) would be too fragmented in larger islands, while larger grids (e.g., 20 km) risked averaging out important spatial variation, especially in smaller islands.

Separate PAMs were generated for (1) combined dataset, (2) preserved-specimen records, (3) citizen-science records, and (4) material-citation records to determine if observed species richness patterns differ among datasets. For each dataset, the unique number of species was calculated per 10-km grid cell, and the results were visualized as spatial maps using QGIS 3.4 Madeira [56]. The resulting maps of herpetofaunal species richness were overlaid with shapefiles of PCAs to assess spatial overlap between observed species richness and PCA coverage area. Geospatial data for protected and conserved areas, as well as key biodiversity areas [63], were sourced from the National Integrated Protected Area System dataset provided by the Department of Environment and Natural Resources – Biodiversity Management Bureau (DENR-BMB). National boundary shapefile was obtained from the National Mapping and Resource Information Authority (NAMRIA). All data were accessed via the Geoportal Philippines, the central repository for official Philippine geospatial information (https://www.geoportal.gov.ph/).

To assess whether species richness classes were evenly represented across grid cells, a chi-square goodness-of-fit test was conducted using the chisq.test function in R version 4.3.1 [64]. Richness values were grouped into nine discrete classes based on the observed range, with 90 as the maximum per grid cell (i.e., 0, 1–10, 11–20, …, 81–90). The observed frequency of cells in each class was compared to a uniform distribution, under the assumption that all classes were equally likely. Expected counts were computed by dividing the total number of grid cells by the number of richness classes. All expected frequencies exceeded 5, meeting assumptions of the chi-square test.

### Predictors of observed species richness across PCAs

Species richness is shaped by a complex interplay of environmental and evolutionary processes. However, the richness values analyzed here reflect observed species counts derived from available occurrence records. Accordingly, our analysis focuses on variation in observed species richness across PCAs and the influence of spatial attributes, rather than attempting to infer the underlying processes that generate biodiversity per se. Specifically, we examined whether area, topographic relief, and sampling effort help account for variation in observed richness, and whether these relationships differ between PCA types. As a supporting analysis, we also tested whether sampling effort is predicted by these same spatial attributes of PCAs, and whether these relationships vary between PCA types. Calculations for each variable are outlined below, followed by a description of our modeling framework.

To quantify species occurrence records and observed species richness within PCAs, we spatially intersected occurrence points with PCA boundaries using “Join Attributes by Location” tool in QGIS 3.4 Madeira [56] (input: occurrence points; join layer: PCA polygons; predicate: intersects). The resulting dataset was exported as a CSV file and processed in R statistical software [64], where occurrence records without PCA matches were excluded. Data were grouped by PCA to calculate the number of occurrence records and unique species per PCA. PCAs without intersecting occurrences were retained with zero counts. Polygon area (in km²) was calculated using expanse function in the R package terra [65]. Only terrestrial PCAs were included in the analysis, comprising 207 protected and conserved areas and 109 key biodiversity areas.

Topographic relief for each PCA was extracted using a Digital Elevation Model (DEM) of the Philippines downloaded from the CGIAR Consortium for Spatial Information (https://srtm.csi.cgiar.org/). This DEM, provided in Arc/Info Grid format, was clipped to the spatial extents of each PCA. Clipping was performed using QGIS’ ‘Clip Raster by Mask Layer’ tool, with DEM as the input raster and PCA polygon shapefiles as the mask layer. The resulting clipped DEM layers contained elevation data exclusively within each PCA boundary. Subsequently, minimum and maximum elevation values were calculated for each PCA polygon using the ‘Zonal Statistics’ tool in QGIS. The difference between these values was used to quantify topographic relief (in meters) of each PCA.

To test whether area, topographic relief, and sampling effort explain variations in observed species richness across PCAs, we fitted generalized linear mixed-effects models (GLMMs) with a negative binomial error distribution to account for overdispersion (dispersion parameter = 0.7466; AIC = 1351.5), which improved model fit relative to a Poisson model (dispersion parameter = 2.1781; AIC = 1601.6). Fixed effects included total area (km²), occurrence density (number of records per km²), topographic relief (m), PCA type, all pairwise interactions among spatial predictors, and interactions between PCA type and each spatial predictor. Area and topographic relief were used as proxies for landscape features, based on the hypothesis that larger PCAs and those spanning broader elevational gradients — thus encompassing more heterogeneous habitats — support higher species richness [66,67]. Occurrence density was included as a proxy for sampling effort, while acknowledging that it may also reflect underlying biological variation in abundance and detectability. As such, it represents a composite measure rather than a purely independent estimate of survey effort, and results involving this variable are interpreted with this limitation in mind. Predictor variables were transformed to improve model fit: area and occurrence density were log□□-transformed due to strong right skew, and topographic relief was square-root-transformed due to moderate skew. All predictors were then mean-centered and scaled to facilitate convergence and allow direct comparisons of effect sizes.

PCA type interactions were included to test whether the relationships between species richness and each spatial predictor differ between protected and conserved areas (managed) and key biodiversity areas (unmanaged). This framework allows comparison of effect sizes and formally evaluates whether the strength of these relationships varies between designation types. To account for potential spatial non-independence among PCAs within biogeographic regions (i.e., PCAs share more similarity in species composition by virtue of historical biogeographic structuring), we included the Pleistocene Aggregate Island Complex (PAIC; see [27]) as a random intercept in all models. This structure reflects the analytical scale of inference, which is defined at the level of heterogenous PCA polygons rather than individual occurrence points. Model selection supported retention of the hypothesized full interaction structure and the random intercept, which had the lowest AICc among candidate models (AICc=1352.75, ΔAICc=0) and was therefore used for inference (S1 Table). Residual diagnostics on the best-selected model indicated minor deviations for occurrence density (S1 Fig), likely reflecting zero inflation from undersampled sites rather than model misspecification. Tests of overdispersion and residual uniformity indicated no violations of model assumptions (S1 Fig).

To further facilitate interpretation, we modeled the number of occurrence points separately to assess the extent to which variation in sampling may influence observed species richness patterns. Total occurrence records per PCA were modeled as a function of PCA type and its interaction with area, topographic relief, and the area × topographic relief interaction, with PAIC again included as a random effect. A negative binomial error distribution was used to account for overdispersion (dispersion parameter = 1.574; AIC = 2561.9), which substantially improved model fit relative to a Poisson model (dispersion = 660.4; AIC = 102443). Residual diagnostics indicated no major violations of model assumptions, although some quantile deviations were observed for each predictor, likely reflecting increased variance at large PCAs where fewer observations exist (S2 Fig). This sampling-effort model provide statistical evidence for whether PCAs with lower observed species richness may be undersampled.

Generalized linear models were fitted using the R package glmmTMB [68]. Overdispersion was initially assessed using a custom R function that calculates the ratio of the sum of squared Pearson residuals to residual degrees of freedom, with values >1.5 considered indicative of overdispersion requiring correction [69]. Model diagnostics for final models were conducted using the DHARMa package [70]. Simulated residuals were generated with the simulateResiduals function, and goodness-of-fit tests on scaled residuals were done using testZeroInflation, testUniformity, and testDispersion functions. Residuals were also plotted against each fixed predictor using plotResiduals to check for non-random patterns. Variance inflation factors (VIFs) were calculated using the R package performance [71]. In the interaction model, several interaction terms showed moderately elevated VIF values (>5), which is expected due to shared variance introduced by interaction structures and does not necessarily indicate problematic collinearity among the primary predictors. To evaluate collinearity among main effects independently of interaction terms, we fitted a reduced model containing only additive predictors. All main-effect predictors in this model showed low multicollinearity (VIFs < 5). All sampling effort models, with occurrence points as the response variable, showed low VIF values (< 5.2). All diagnostic outputs and residual plots are provided in S1–S2 Figs.

As a complement to model-based inference, bivariate relationships were visualized using negative binomial generalized linear models (log link) with 95% confidence intervals, fitted separately by PCA type using the geom_smooth function in the R package ggplot2 [72]. Predictor variables were transformed to match model specifications, and point colors were used to reflect the values of interacting variables with significant effects, unless otherwise stated, aiding interpretation of interaction patterns. All statistical analyses were conducted in R version 4.3.1 [64] and documented in Supplementary Material.

## Results

### Spatial distribution of herpetofaunal diversity in the Philippines

Spatial distribution of observed herpetofaunal species richness in the Philippines is highly non-uniform and fragmented, with unequal national-scale coverage that varies considerably across islands (Fig 1A). The distribution of species richness classes deviated significantly from a uniform expectation (χ²(8) = 15,843, *p* < 0.001), indicating uneven representation of richness classes in the observed dataset. Some richness classes were overrepresented, while others were underrepresented relative to expectations if all classes were equally likely. Nationally, only ∼2% (n = 87) of the 10-km² grids have more than 41 observed species—approximately 50% of maximum richness recorded in any grid—while 31% (n = 1,370) have between 1 and 40 species, and 66.7% (n = 2,914) have zero recorded species.

**Fig 1.**
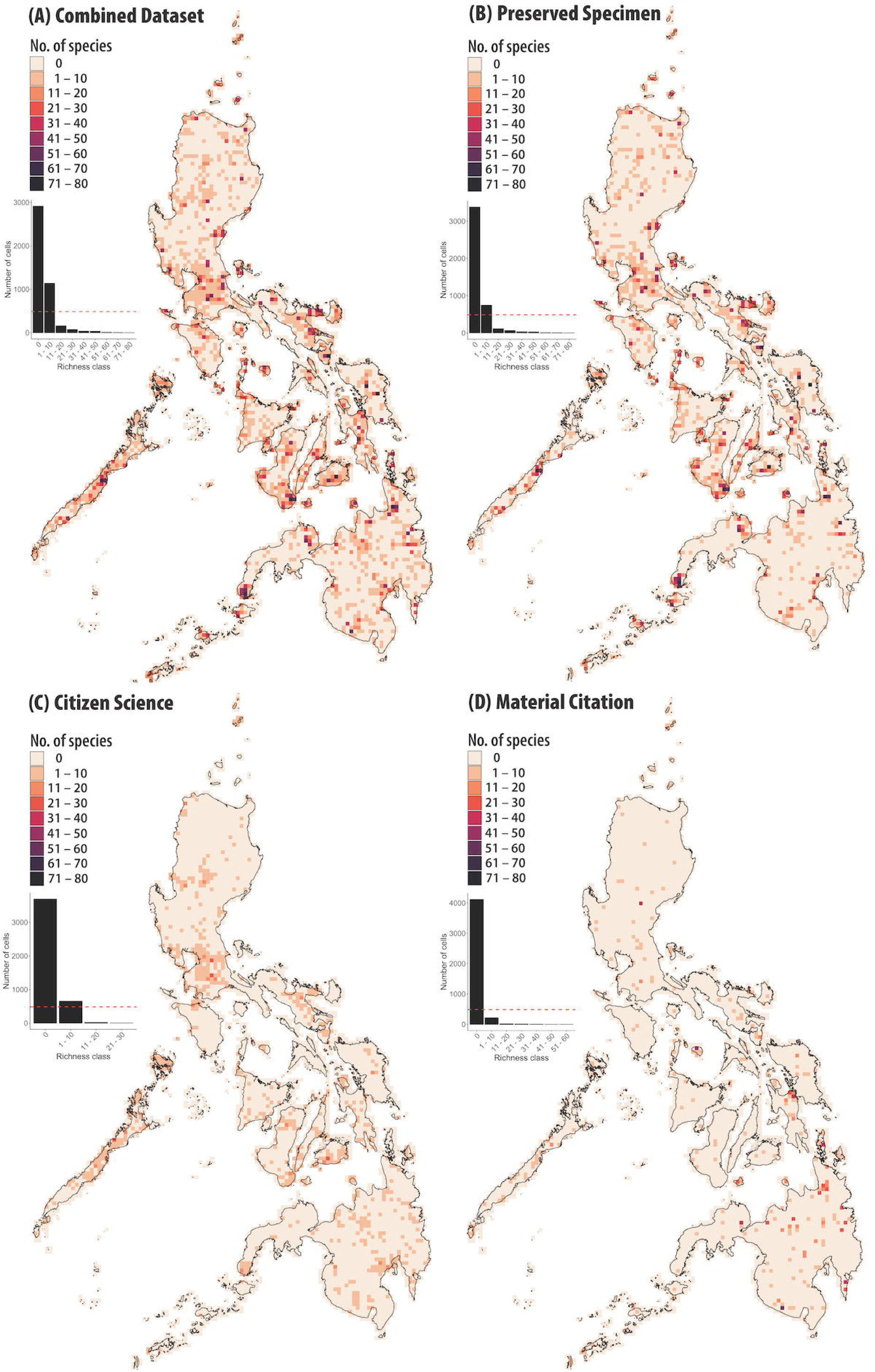
Spatial distribution of observed amphibian and squamate reptile species richness for (A) combined datasets; (B) preserved-specimen records; (C) citizen-science records; and (D) material-citation records. Species richness is calculated per 10-km² grid cell across the Philippines. Inset bar graph shows frequency distribution of grid cells across species richness classes for each dataset. Dashed red line indicates expected frequency for each class under a uniform distribution, as used in chi-square goodness-of-fit tests. Administrative boundary and biodiversity area shapefiles were sourced from NAMRIA and the DENR-BMB via Geoportal Philippines (https://www.geoportal.gov.ph/).

Different data sources provide complementary but uneven contributions to observed herpetofaunal species richness patterns. The majority of occurrence records came from preserved specimens (n = 70,112), followed by citizen science contributions (n = 3,735) and material citations (n = 2,344), while 37 points were unclassified. Both preserved-specimen and citizen-science data represent individual occurrences, whereas material citations typically reflect species-level records, as published species inventory studies mostly report presence by species rather than by individuals. All data sources significantly deviated from a uniform expectation (χ²(8) = 20,186–30,610, *p* < 0.001). Because preserved specimens account for 92% of all records, broad-scale spatial richness patterns are largely influenced by specimen-based sampling (Fig 1B). Citizen science records generally contribute 1–10 species per grid, with a few grids reaching up to 30 species, and tend to overlap with already sampled regions (Fig 1C). Material citation data are more spatially restricted, consistent with the targeted survey designs and site-based biotic inventories (Fig 1D).

Philippine conservation-relevant areas (PCAs) show variable representation in the occurrence dataset, although observed species richness remains uneven across sites (Fig 2). Of the 76,228 curated occurrence records, 21,210 (27.8%) fell within protected and conserved areas and 23,058 (30.2%) within key biodiversity areas—higher proportions than expected based on the total land area covered by these designations (12–15% for protected and conserved areas and 6.72% for key biodiversity areas). When overlaid with richness patterns, some high-richness grid cells occur within PCA boundaries, while many large PCAs include grid cells with few or no records. Notably, 52.7% of 207 protected and conserved areas and 20.2% of 109 key biodiversity areas contain grid cells with zero species records, and an additional 22.4% and 29.4%, respectively, contain only up to 10 recorded species. At the same time, several high-richness grid cells (>40 recorded species) occur outside protected and conserved areas but within forested landscapes, some of which partially overlap with key biodiversity areas (S3 Fig).

**Fig 2.**
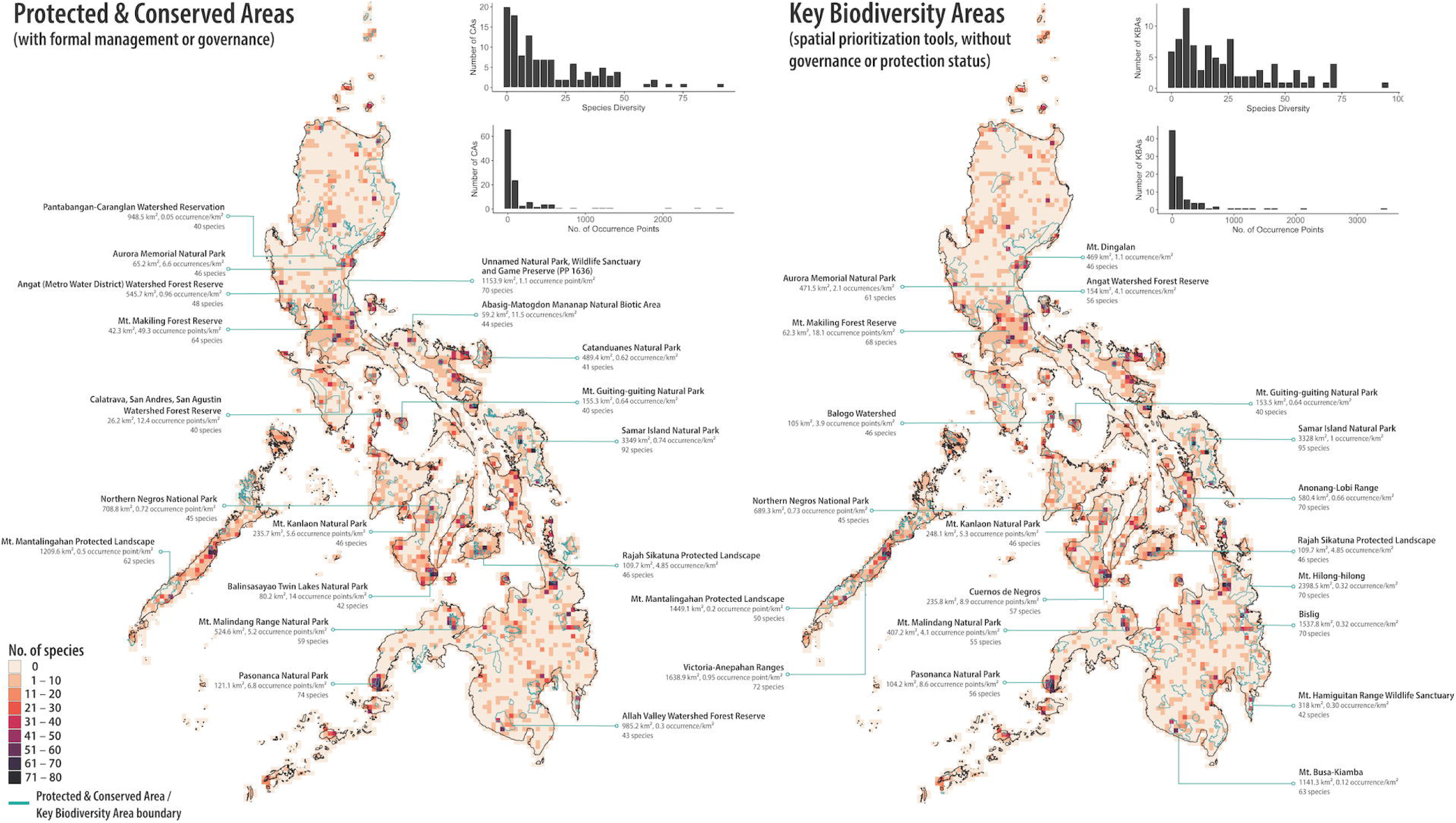
Relative locations of well-sampled (≥40 species) conservation-relevant areas across the Philippines (PCAs). Total area, occurrence point density, and number of species are included for each area. Base map shows spatial distribution of observed amphibian and squamate reptile richness patterns for combined data sources. Inset histogram shows frequency distribution of PCAs by species richness and number of occurrence records. Administrative boundary and biodiversity area shapefiles were sourced from NAMRIA and the DENR-BMB via Geoportal Philippines (https://www.geoportal.gov.ph/).

Well-surveyed sites (>40 recorded species) make up 8.8% of protected and conserved areas and 18.3% of key biodiversity areas (Fig 2). Sites with the highest recorded richness (>60 species) include Samar Island Natural Park, Pasonanca Natural Park, Mt. Malindang Natural Park, Mt. Makiling Forest Reserve, and PP1636 (unnamed wildlife sanctuary in southcentral Luzon). These areas are among the most intensively sampled in available datasets, with records distributed across public repositories such as GBIF, although additional data may remain unpublished in peer-reviewed literature (but see [74,75]). Other relatively well-documented areas combine strong specimen representation in collections with published data: Cuernos de Negros [76,77], Aurora Memorial National Park [78,79], Mt. Busa–Kimba [61,79], Mt. Guiting-Guiting [80,81], Mt. Hilong-hilong [60,82], Pantabangan-Caranglan Watershed Reservation [83], and Victoria-Anepahan Ranges [84]. The ≥40 species threshold is used here only as a descriptive reference and should not be interpreted as a biologically meaningful cutoff for “high diversity,” as observed richness is influenced by sampling effort, detectability, and environmental heterogeneity.

### Predictors of observed species richness across PCAs

Model selection based on AICc supported the hypothesized interaction model as the best-supported candidate model (S1 Table), although several reduced models also received substantial support (ΔAICc < 2). There was insufficient evidence of a difference in baseline species richness between key biodiversity areas (reference category) and protected and conserved areas (Table 1). In key biodiversity areas, observed species richness increased with total area, topographic relief, and occurrence density; larger key biodiversity areas supported more species (β_area = 1.2412 ± 0.1142 SE, *z* = 10.82, *p* < 0.001; Fig 3A), richness increased strongly with occurrence density (β_occurrence density = 1.5402 ± 0.0862, *z* = 17.875, *p* < 0.001; Fig 3B), and areas encompassing greater topographic relief were richer (β_topographic relief = 0.3265 ± 0.1098, *z* = 2.973, *p* = 0.003; Fig 3C). Two-way interactions among predictors indicated non-additive effects of spatial attributes. Specifically, the positive effect of area on richness weakened in key biodiversity areas with greater topographic relief (area × topographic relief: β = –0.2338 ± 0.0643, *z* = –3.637, *p* < 0.01), which may suggest that the species–area relationship is less steep in more topographically heterogeneous landscapes. Likewise, the positive effect of occurrence density on richness decreased slightly as topographic relief increased (occurrence density × topographic relief: β = –0.2031 ± 0.0830, *z* = –2.446, *p* = 0.014).

**Fig 3.**
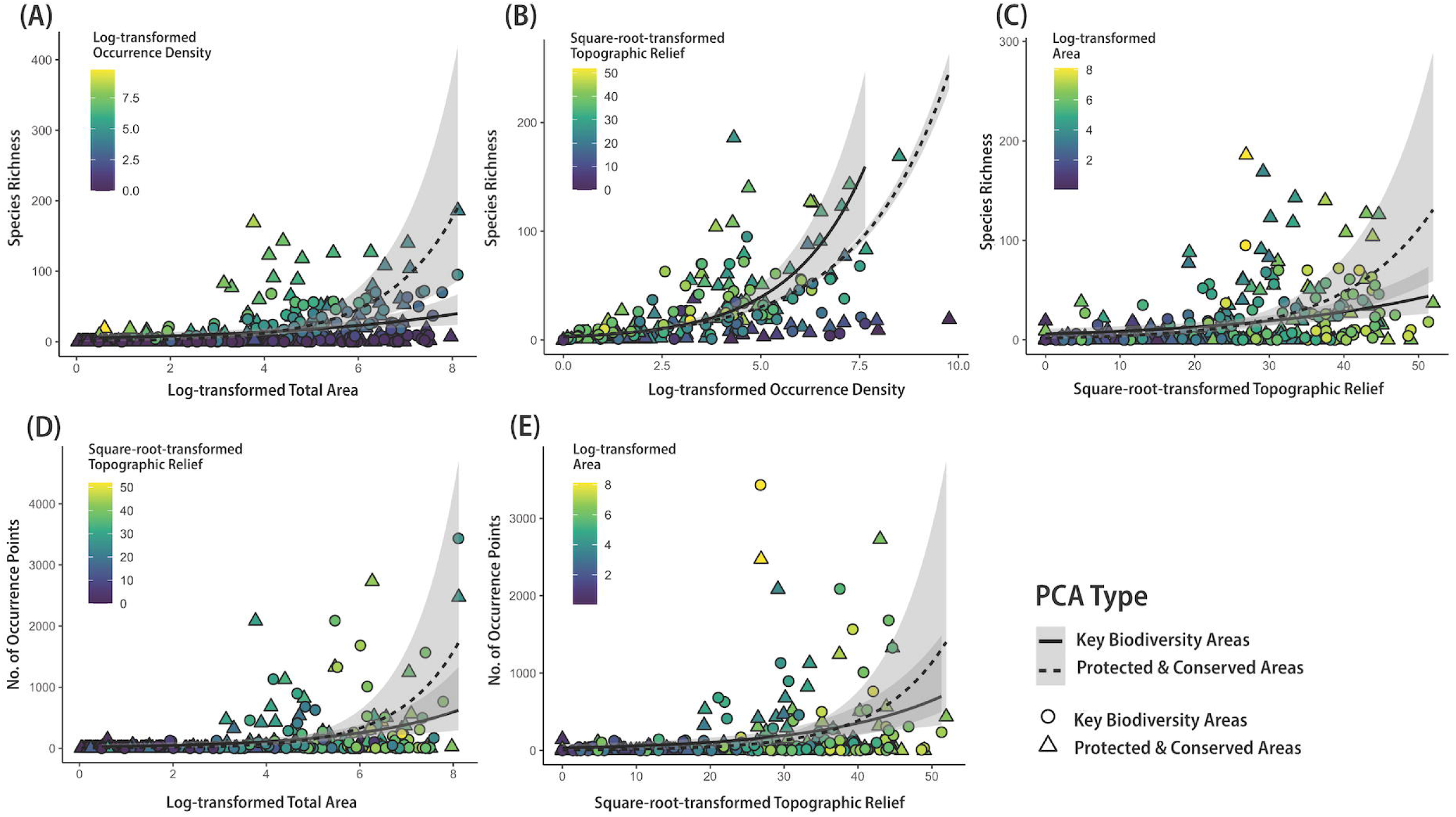
Bivariate relationships between species richness and total area, occurrence density, and topographic relief; and between sampling effort and area and topographic relief across Philippine conservation-relevant areas (PCAs). Lines represent negative binomial generalized linear model fits (log link) estimated separately by PCA type, with grey bands indicating 95% confidence intervals. Points represent individual PCAs; colors denote interacting predictors.

**Table 1.**
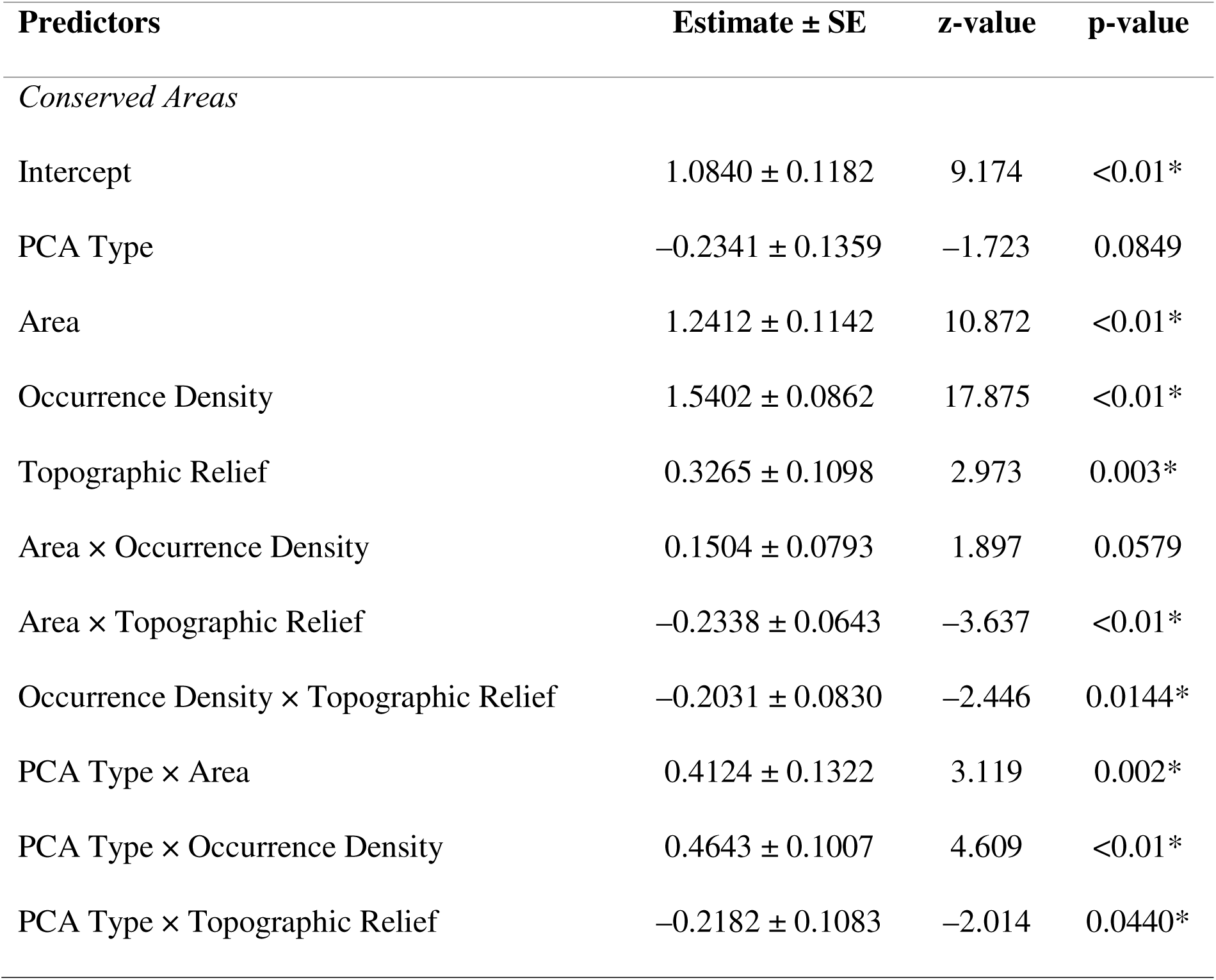
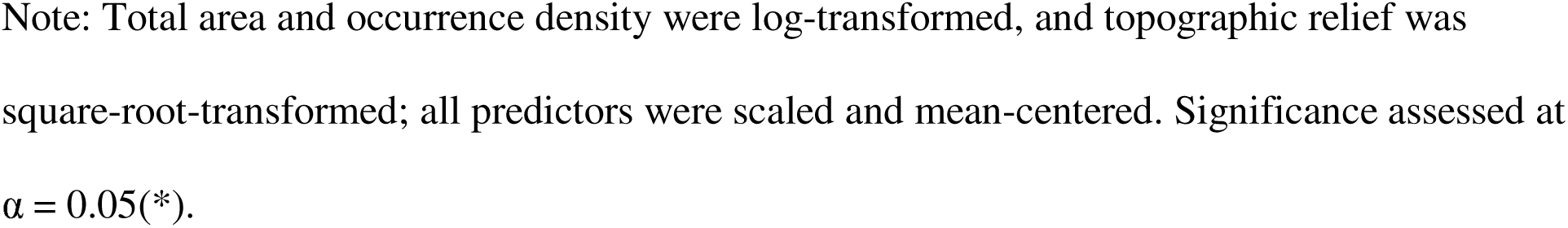
Coefficient estimates from a generalized linear mixed-effects model (negative binomial), in which species richness was modeled as a function of area, topographic relief, and occurrence density. The model included all pairwise interactions among these predictors, as well as interactions between each predictor and Philippine conservation-relevant area (PCA) type (reference category: key biodiversity area). Biogeographic

| Predictors | Estimate ± SE | z-value | p-value |
| --- | --- | --- | --- |
| *Conserved Areas* |  |  |  |
| Intercept | 1.0840 ± 0.1182 | 9.174 | <0.01* |
| PCA Type | -0.2341 ± 0.1359 | -1.723 | 0.0849 |
| Area | 1.2412 ± 0.1142 | 10.872 | <0.01* |
| Occurrence Density | 1.5402 ± 0.0862 | 17.875 | <0.01* |
| Topographic Relief | 0.3265 ± 0.1098 | 2.973 | 0.003* |
| Area × Occurrence Density | 0.1504 ± 0.0793 | 1.897 | 0.0579 |
| Area × Topographic Relief | -0.2338 ± 0.0643 | -3.637 | <0.01* |
| Occurrence Density × Topographic Relief | -0.2031 ± 0.0830 | -2.446 | 0.0144* |
| PCA Type × Area | 0.4124 ± 0.1322 | 3.119 | 0.002* |
| PCA Type × Occurrence Density | 0.4643 ± 0.1007 | 4.609 | <0.01* |
| PCA Type × Topographic Relief | -0.2182 ± 0.1083 | -2.014 | 0.0440* |

Note: Total area and occurrence density were log-transformed, and topographic relief was square-root-transformed; all predictors were scaled and mean-centered. Significance assessed at $\alpha = 0.05(*)$ .

Relative to key biodiversity areas, protected and conserved areas exhibited significantly steeper species–area relationships (PCA type × area: β = 0.4124 ± 0.1322, *z* = 3.119, *p* = 0.002; Fig 3A) and stronger positive effects of occurrence density (PCA type × occurrence density: β = 0.4643 ± 0.1007, *z* = 4.609, *p* < 0.01; Fig 3b), whereas the effect of topographic relief on species richness was weaker in protected and conserved areas (PCA type × topographic relief: β = – 0.2182 ± 0.1083, *z* = –2.014, *p* = 0.044; Fig 3C). The random intercept for biogeographic subregion (PAIC) explained negligible variance in richness ( variance ≈ 6.87 × 10□¹□; SD ≈ 2.62 × 10□□), which may suggest minimal influence of biogeographic clustering on observed richness.

Only occurrence density × topographic relief and area × topographic relief interactions were statistically significant.

To evaluate potential influence of sampling effort, total occurrence records per PCA were modeled as a function of area, topographic relief, and PCA type. We found no strong evidence for differences in baseline number of occurrence points between key biodiversity areas and protected and conserved areas (Table 2). In key biodiversity areas, number of occurrence points showed marginal positive relationships with area (β = 0.7267 ± 0.3782, *z* = 1.922, *p* = 0.055; Fig 3D) and topographic relief (β = 0.7181 ± 0.3854, *z* = 1.863, *p* = 0.062; Fig 3E), and none of the interactions with PCA type were significant. The PAIC random intercept accounted for moderate variance in sampling (variance = 0.46; SD = 0.68), potentially reflecting geographic clustering in sampling effort and/or underlying variation in biodiversity across biogeographic regions. Together, these results suggest that differences between PCA types are unlikely to be driven by strong systematic differences in sampling effort, although some contribution of sampling heterogeneity cannot be fully excluded. This interpretation should be considered cautiously because occurrence records represent an imperfect proxy for sampling effort and may also reflect underlying biological signal.

**Table 2.** Coefficient estimates from a generalized linear mixed-effects model (negative binomial) in which number of occurrence points was modeled as a function of area, topographic relief, area × topographic relief, and their interactions with Philippine conservation-relevant area (PCA) type (reference category: key biodiversity area). Biogeographic subregion was included as a random intercept.

| Predictors | Estimate ± SE | z-value | p-value |
| --- | --- | --- | --- |
| Intercept | 5.0046 ± 0.4661 | 10.737 | < 0.01 * |
| PCA Type | -0.5019 ± 0.4034 | -1.244 | 0.2134 |
| Area | 0.7267 ± 0.3782 | 1.922 | 0.0547 |
| Topographic Relief | 0.7181 ± 0.3854 | 1.863 | 0.0624 |
| Area × Topographic Relief | -0.5365 ± 0.3187 | -1.683 | 0.0923 |
| PCA Type × Area | 0.1607 ± 0.4746 | 0.339 | 0.7350 |
| PCA Type × Topographic Relief | 0.3017 ± 0.5100 | 0.592 | 0.5541 |
| PCA Type × Area × Topographic Relief | 0.2050 ± 0.3862 | 0.531 | 0.5956 |

Note: Total area was log-transformed and topographic relief was square-root-transformed; continuous predictors were scaled and mean-centered. Significance assessed at $\alpha = 0.05(*)$ .

## Discussion

### Broad-scale geographic heterogeneity in biodiversity data

Our spatial mapping of observed species richness of amphibians and squamate reptiles revealed stark patterns of sampling biases across the Philippines. More than two-thirds of grid cells contain either zero or only 1–10 documented species, while comparatively fewer cells (∼2%) exhibit higher species richness levels. This pronounced disparity likely reflects persistent spatial biases in biodiversity knowledge. Observed species richness patterns likely reflect, in part, where sampling has been more intensive, potentially limiting the extent to which observed richness can be interpreted independently of sampling structure and underlying ecological or biogeographic variation. Nevertheless, this pattern is persistent across many biodiversity-rich areas globally [17,20,21,85]. Addressing these biases is critical not only for improving knowledge of Philippine biodiversity but also for ensuring that conservation policy, priority-setting, and management decisions rest on a more complete and representative knowledge base.

A recent comprehensive review of Philippine herpetology revealed substantial geographic gaps in field sampling across the archipelago (see [25] for a detailed discussion). Our findings broadly align with these observations but offer additional fine-scale information by integrating curated species point data and grid-based species richness estimates. This approach not only identifies where sampling has occurred but also quantifies its intensity, suggesting that higher species richness values are likely associated with well-sampled sites (i.e., more occurrence points). Notably, we found that a disproportionate share of documented herpetofaunal diversity is concentrated in a few intensively surveyed areas—primarily in Luzon (the largest island) and in the West Visayas, which comprise much of the central islands—where collection effort has historically been high [25,39]. However, many records from the West Visayas date to pre-2000 surveys and warrant resampling [25]. High-resolution data of this kind are valuable for characterizing spatial patterns of sampling effort, thereby guiding resources and research efforts towards poorly explored regions. Furthermore, by incorporating species point data from different sources, we recovered diversity data in areas that would otherwise appear as knowledge gaps, such as the Sulu Archipelago in the south [25]; although sampling effort is still disproportionately lower in this region. As such, given that biodiversity data in the Philippines are fragmented, reliance on selective or incomplete datasets for *any* synthesis studies can introduce substantial bias [86].

Philippine conservation-relevant areas (PCAs) are relatively well-sampled in proportion to their total area, yet herpetofaunal knowledge across them remains limited and uneven. Despite these gains, many PCAs still lack species records altogether or may have data that are not accessible in digital form. Where species data are lacking in PCAs, they are however available outside delineated boundaries, which may reflect persistent logistical and institutional constraints associated with sampling within PCAs, including permitting and slow bureaucratic processes (particularly for legislated protected areas), access, and security issues [25,60,87] (see Fig 4). Although well-surveyed sites include some of the country’s known PCAs, vast portions of other mountain ranges and many smaller islands remain poorly sampled or lack species data that are readily accessible for broader scientific and conservation use. Expanding targeted field-based surveys beyond established PCAs—and ensuring that resulting data are made publicly and digitally available for mobilization—remains urgently needed to support conservation initiatives grounded in more representative and spatially comprehensive biodiversity data [21,88].

**Fig 4.**
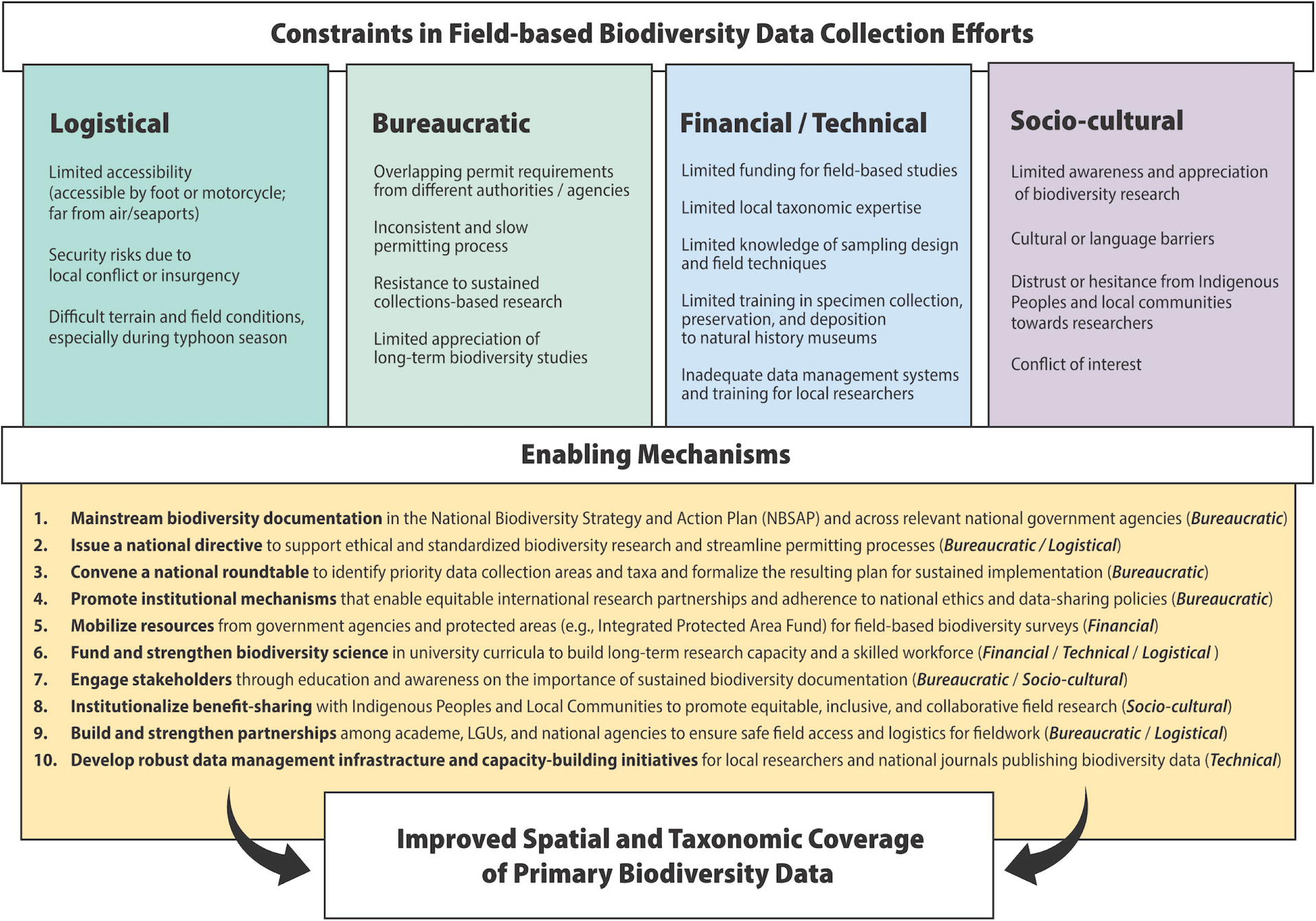
Constraints in field-based biodiversity data collection and corresponding enabling mechanisms to improve primary biodiversity data coverage. Each mechanism primarily addresses one or more constraint types.

### Predictor effects vary across conservation-relevant areas

Our unified modeling framework provides no strong statistical evidence that baseline species richness differs between types of conservation-relevant areas in the Philippines (protected and conserved areas vs. key biodiversity areas) after accounting for area, sampling effort, and topographic relief. However, the relationships between species richness and spatial predictors vary between designation types, as indicated by significant interaction terms. In both PCA types, area, occurrence density, and topographic relief were positively associated with observed richness. The strength of the species–area relationship and the effect of occurrence density were greater in protected and conserved areas, whereas the effect of topographic relief was relatively stronger in key biodiversity areas. Importantly, these differences are unlikely to be explained by strong systematic variation in sampling effort between designation types. Our sampling effort model found no clear difference in baseline number of occurrence points between PCA types, and no evidence that PCA type modifies the relationships between number of occurrence points and either area or topographic relief. However, because occurrence records may also reflect underlying biological variation in abundance or detectability, biological-related effects cannot be fully excluded. Overall, the stronger richness–area and richness–sampling relationships observed in protected and conserved areas are unlikely to be driven solely by uneven survey effort.

The stronger elevational effect in key biodiversity areas suggests that observed species richness shows a steeper association with elevation in these areas compared to protected and conserved areas. One possible ecological interpretation of this pattern is that key biodiversity areas are planning tools that target areas of high ecological importance, such as concentrations of endemic or range-restricted taxa, many of which occur in topographically complex regions [89]. As such, their boundary delineations may encompass broader gradients of topographic relief and associated habitat diversity along elevation. In contrast, protected and conserved area designation involves additional socio-political and land-use considerations that may not always prioritize elevational heterogeneity [90–92]. In the Philippines, nationally legislated protected areas are designated based not only on biodiversity value but also on socio-cultural importance and provision of key ecosystem services, including regional water security, which may be politically influenced [93,94]. In addition, our definition of protected and conserved areas includes other effective area-based conservation measures, such as Indigenous and Community Conserved Areas and locally managed conservation areas, whose establishment is often motivated by social and governance considerations beyond ecological criteria [92].

Between both designation types, the positive effect of topographic relief weakens as area and sampling intensity increase, as indicated by significant negative interaction terms. This pattern differs from the general ecological expectation that large, topographically complex areas tend to support higher diversity due to habitat heterogeneity [17,66,67,95]. One plausible explanation is that elevational heterogeneity may promote species turnover at smaller spatial extents; however, as area increases, portions of this gradient are increasingly represented within sampling units, which may lead to partial redundancy in observed species composition (see scale effects on elevational species richness [96]). This pattern is also consistent with the possibility that sampling coverage becomes more spatially saturated in larger units, although the relative contributions of ecological structure and sampling completeness cannot be fully separated in the present dataset. Similarly, in more intensively sampled PCAs, additional survey effort may disproportionately detect widespread or low-elevation taxa rather than reveal the full complement of high-elevation specialists, many of which are rarely documented and taxonomically undescribed [42,60,62,75,81,97].

Sampling constraints may also contribute to this pattern. Mountainous regions in the Philippines are frequently remote, difficult to access, and logistically expensive to survey [97] (see Fig 4). Even where PCAs encompass wide topographic relief, field effort may be concentrated in more accessible foothill zones, potentially leaving high-elevation habitats underrepresented [25]. This limitation has been highlighted in recent assessments of Philippine herpetological sampling [25]. As a result, observed richness may underrepresent the contribution of topographically complex systems, potentially biasing biodiversity knowledge toward more accessible montane zones despite their well-documented ecological importance and high levels of local endemism [38, 98]. Together, these findings suggest that large, mountainous PCAs do not necessarily translate into proportionally higher recorded richness — not because they lack diversity, but because the effects of elevational heterogeneity may be scale-dependent and partly obscured by uneven sampling coverage.

Biodiversity data limitations are not confined to large mountain ranges. Smaller conservation areas, particularly those in island environments, also suffer from substantial knowledge gaps [51]. Although this pattern was not statistically prominent in our model-based inference, it was evident in our data visualizations (e.g., small-sized, low-topographic-relief PCAs in Fig 3). Many small island conservation areas in the Philippines remain poorly sampled despite yielding newly discovered species and harboring unique, range-restricted taxa [99–107]. These island systems face disproportionate threats from climate-driven sea-level rise, extreme weather events, invasive species, and habitat loss [108], yet lack the baseline biodiversity data for adaptive management. Targeted surveys are urgently needed in these areas to document species presence and build taxonomic and ecological knowledge base required for effective conservation in islands [109]. Without action, island ecosystems risk becoming critical blind spots in the country’s conservation efforts.

### Data gaps and biases persist, but diverse sources can address them

The majority of digitally accessible knowledge of Philippine herpetofauna comes from natural history collections. These collections represent over a century of fieldwork that has shaped taxonomic and systematics knowledge, increasing herpetofaunal diversity estimates in the country [25,39]. As such, specimen-associated data serve as a vital resource for characterizing biodiversity patterns [5,110]. They also support large-scale ecological studies, which depend on spatially referenced records to examine broad biodiversity trends and inform conservation planning [16,31,111]. Although many regions remain underrepresented in biodiversity data, it is noteworthy that areas with limited contemporary surveys—as mentioned earlier, Sulu Archipelago, where fieldwork has been limited [25]—still hold valuable records preserved through historical collections, many of which are the only verifiable species records for this biogeographic subregion.

In areas where specimens are lacking, citizen science provides a valuable complementary data stream that can help reduce some geographic gaps in biodiversity records. Online biodiversity platforms [112–114] and social media [115–118] have become particularly useful where museum records are absent or limited. Although still emerging in Philippine herpetology (but see [118–120]), citizen-science contributions are already well established in other taxonomic groups. For example, active birding communities regularly contribute to platforms like eBird [121] and collaborate with researchers and biodiversity managers to inform site-level conservation efforts in the Philippines [94]. Another notable initiative is Co’s Digital Flora of the Philippines [122], where citizen scientists and taxonomists work together to maintain a real-time overview of Philippine flora. At least 58% of the country’s vascular plant species have been photo-documented, many with associated geographic coordinates [41]. These examples show that citizen-science contributions to observed diversity may be more pronounced in taxonomic groups with active, organized communities driving such efforts. Despite challenges related to data quality and metadata completeness, carefully curated citizen-science records can enhance biodiversity data coverage, particularly in remote or poorly sampled regions [112,123].

Another important yet often overlooked source of biodiversity data comes from formal surveys that do not involve specimen collection and eventual deposition in natural history museums [61,83,124–126]. And if there are such records, they are often not digitized and published in publicly accessible domains, especially when deposited in university-based museums. Many in-country scientists conduct fieldwork that reports valuable records published in peer-reviewed journals, yet these data are rarely archived in open-access databases such as GBIF [127]. Such studies frequently document species in poorly sampled areas in the Philippines, providing crucial complementary information [25]. Although valuable for expanding species distribution knowledge, many of these studies lack specific geographic coordinates of areas sampled, limiting their use for geospatial research; thus, submission of spatial occurrences to online databases like GBIF is highly encouraged [36]. In addition, a wealth of biodiversity data remains locked in grey literature—government reports, university theses, and project documents—that are not digitally accessible but are often used in site-level management. Incorporating these sources into public repositories and ensuring they are properly curated would improve national biodiversity coverage [128].

Disparate biodiversity data from different sources underscore the need for standardized archiving practices [129–131] and greater adherence to best-practice guidelines in dealing with big data [20,132]. Although these varied data streams help fill distributional gaps, they often differ in quality, accessibility, and curation. These differences are especially true for non-specimen-based data, which typically lack the standardized metadata (Darwin Core format) associated with museum specimens. To maximize their scientific utility, we recommend that non-specimen-based records be accompanied by metadata, including GPS coordinates (including uncertainty), observation date and time, natural history notes, among others. Robust metadata not only enhance credibility and utility of individual records but also facilitates their integration into broader ecological, biogeographic, and conservation research [16].

### Scaling biodiversity documentation to meet conservation targets

The Kunming-Montreal Global Biodiversity Framework, adopted in 2022, sets an ambitious goal: to protect 30% of the world’s terrestrial and marine ecosystems by 2030, building on the earlier Aichi Target 11 [133]. Achieving this target requires more than simply expanding protected area coverage; it also demands effective management that delivers measurable benefits for biodiversity [30]. These outcomes depend on robust, spatially representative biodiversity data, particularly within conservation-relevant areas in the Philippines [34,134,135]. However, our results show that persistent data gaps and biases, especially in large mountainous and island conservation areas, continue to shape observed biodiversity data patterns. Although many PCAs with apparent data gaps may in fact hold biodiversity data, their lack of digital accessibility limits inclusion in national-scale assessments of sampling effort. This limitation hinders resource allocation for improved documentation, reduces opportunities for external vetting and expert curation of data, and impedes integration into broader datasets that inform national conservation planning [21,31,130,136].

Addressing the challenge of insufficient biodiversity documentation requires developing and implementing enabling mechanisms to overcome constraints in site-level primary data collection, which comprises the majority—if not all—of available biodiversity data (Fig 4). This includes prioritizing biodiversity documentation within national conservation policies and planning frameworks (e.g., Philippine Biodiversity Strategy and Action Plan, PBSAP 2024–2040) to strengthen institutional support for field-based activities, especially within PCAs [23]. For example, streamlining research permitting processes, fully aligned with Indigenous Peoples’ rights and national access and benefit-sharing policies, can reduce bureaucratic barriers to ethical and standardized data collection [36,137]. Engaging Indigenous Peoples and local communities as collaborators, and supporting culturally appropriate awareness and communication initiatives, can further foster equitable partnerships and mutual knowledge exchange [138]. Financial mechanisms within national legislated Protected Areas, such as the Integrated Protected Area Fund, can be leveraged to support student-led and locally collaborative field research. Local or subnational government policies and support mechanisms are also critical for ensuring logistical coordination, safe access, and community involvement in fieldwork.

Beyond site-level interventions, targeted national funding for species discovery and taxonomic capacity-building can incentivize early-career researchers to pursue field-based work [137,139]. These efforts should be complemented by investments in biodiversity science within university curricula [140], ensuring students are trained in both theory and practice. Addressing the limited data management capacity among local researchers is also critical for promoting effective data-sharing, enhancing the verifiability and utility of biodiversity studies. Comprehensive training programs and standardized data systems can help overcome this barrier, ensuring the production of high-quality, accessible datasets. In parallel, the increasing diversity of data contributors underscores the need for a streamlined and curated national biodiversity data infrastructure [141]. Such a platform should ensure that biodiversity data, especially those generated from PCAs, are digitally and publicly accessible, enabling diverse stakeholders to collectively contribute to and benefit from biodiversity knowledge. Ultimately, these in-country initiatives must be paired with stronger institutional mechanisms, such as equitable collaboration standards and data-sharing agreements, to ensure that non-monetary benefits from biological resource use are meaningfully shared with local collaborators and inform national policy and conservation planning [142, 143].

Finally, scaling biodiversity documentation beyond existing PCAs can guide the identification of new priority sites for protected area establishment. Our findings show that several well-sampled, high-diversity areas remain outside current PCAs (neither protected/conserved nor delineated as priority areas for conservation), many overlapping with only partially protected key biodiversity areas. This mismatch underscores opportunities to align future protected and conserved area expansion (i.e., protected area, other effective area-based conservation measures) with empirically documented biodiversity patterns, particularly in remaining forested regions. The updated national biodiversity strategy and action plan (PBSAP 2024–2040) targets protecting 24% of terrestrial areas through protected and conserved areas [144], yet only ∼15% are currently designated. Prioritizing well-sampled areas with high observed species richness for protection may help support this goal. Our results, however, only focus on herpetofauna and similar data gaps and biases likely affect many taxonomic groups in the Philippines [24,26] reinforcing the need for broader, inclusive biodiversity documentation. Building a robust and representative biodiversity knowledge base—well-curated and aligned with FAIR (findable, accessible, interoperable, reusable) and CARE (collective benefit, authority to control, responsibility, ethics) principles—will require sustained collaboration among scientists, conservation practitioners, institutions, and critically, with Indigenous Peoples and local communities [16, 143–146]. Strengthening such collective and inclusive efforts is essential for transforming biodiversity knowledge into tangible conservation outcomes and ensuring the Philippines meets its national and global biodiversity commitments.

## Acknowledgements

We thank Marites Bonachita Sanguila, Michael Clores, Michael Cuesta, Aljohn Jay Saavedra, and Tristan Luap Senarillos for numerous discussions and their valuable insights on the challenges of primary biodiversity data collection in the Philippines. We would also like to thank the Editor and two reviewers for their constructive feedback, which greatly improved the manuscript.

## Supporting Information

**Table S1. Top-supported generalized linear mixed-effects models explaining observed herpetofaunal species richness across Philippine conservation-relevant areas (PCAs).** All models included PAIC as a random intercept and were fitted using a negative binomial error distribution. Predictor variables were scaled prior to analysis. “Area” refers to log-transformed PCA area, “Occurrence density” refers to log-transformed occurrence records per 100 km², and “Topographic relief” refers to square-root transformed topographic relief.

**S1 Fig. Model diagnostic results from DHARMa for the species diversity model of Philippine conservation-relevant areas.** (A) Simulated residuals from the fitted model, showing detected quantile deviations. (B) Nonparametric dispersion test. (C) Zero-inflation test. (D) Q–Q plot used to assess deviations from the expected residual distribution, together with tests for uniformity (Kolmogorov–Smirnov), dispersion, and outliers. (E–F) Scaled residuals plotted against predicted values and against each model predictor. Simulation outliers—observations falling outside the range of simulated values—are highlighted as red stars, while red trend lines indicate statistically significant deviations from model expectations. Deviations were primarily associated with the predictor occurrence density, which likely reflects zero inflation arising from undersampled sites rather than true model misspecification. (G) Bivariate relationship between species richness and occurrence density.

**S2 Fig. Model diagnostic results from DHARMa for the sampling effort model of Philippine conservation-relevant areas.** (A) Simulated residuals from the fitted model, showing detected quantile deviations. (B) Nonparametric dispersion test. (C) Zero-inflation test. (D) Q–Q plot used to assess deviations from the expected residual distribution, together with tests for uniformity (Kolmogorov–Smirnov), dispersion, and outliers. (E–F) Scaled residuals plotted against predicted values and against each model predictor. Simulation outliers—observations falling outside the range of simulated values—are highlighted as red stars, while red trend lines indicate statistically significant deviations from model expectations. (G–H) Bivariate relationship between species richness and each model predictor. Deviations were primarily associated with each predictor, likely reflecting increased variance at large PCAs where fewer observations exist.

**S3 Fig. Map of the Philippines,** showing (A) overlap between forest cover and Philippine conservation-relevant areas (PCAs); (B) distribution of observed amphibian and squamate reptile diversity across all combined data sources outside PCA boundaries, highlighting that many well-sampled sites with higher observed species richness lie beyond established conservation-relevant areas. Administrative boundary and biodiversity area shapefiles were sourced from NAMRIA and the DENR-BMB via Geoportal Philippines (https://www.geoportal.gov.ph/).

