## Supplementary Material for "Sampling bias and uneven data coverage shape observed biodiversity patterns in a megadiverse island archipelago hotspot"

**Supporting Information**

**Table S1. Top-supported generalized linear mixed-effects models explaining observed herpetofaunal species richness across Philippine conservation-relevant areas (PCAs).** All models included PAIC as a random intercept and were fitted using a negative binomial error distribution. Predictor variables were scaled prior to analysis. “Area” refers to log-transformed PCA area, “Occurrence density” refers to log-transformed occurrence records per 100 km², and “Topographic relief” refers to square-root transformed elevational breadth.

| Model | Fixed effects structure | df | logLik | AICc | ΔAICc | Weight |
| --- | --- | --- | --- | --- | --- | --- |
| 1 | PCA type + Area + Occurrence density + Topographic relief + Area × Occurrence density + Area × PCA type + Area × Topographic relief + Occurrence density × PCA type + Occurrence density × Topographic relief + PCA type × Topographic relief | 13 | -662.77 | 1352.75 | 0.00 | 0.400 |
| 2 | PCA type + Area + Occurrence density + Topographic relief + Area × PCA type + Area × Topographic relief + Occurrence density × PCA type + Occurrence density × Topographic relief + PCA type × Topographic relief | 12 | -664.56 | 1354.15 | 1.40 | 0.199 |
| 3 | PCA type + Area + Occurrence density + Topographic relief + Area × PCA type + Area × Topographic relief + Occurrence density × PCA type + PCA type × Topographic relief | 11 | -665.84 | 1354.55 | 1.80 | 0.163 |
| 4 | PCA type + Area + Occurrence density + Topographic relief + Area × Occurrence density + Area × PCA type + Area × Topographic relief + Occurrence density × PCA type + Occurrence density × Topographic relief | 12 | -664.83 | 1354.69 | 1.94 | 0.151 |
| 5 | PCA type + Area + Occurrence density + Topographic relief + Area × PCA type + Area × Topographic relief + Occurrence density × PCA type | 10 | -667.54 | 1355.80 | 3.06 | 0.087 |

**S1 Fig. Model diagnostic results from DHARMa for the species diversity model of Philippine conservation-relevant areas.** (A) Simulated residuals from the fitted model, showing detected quantile deviations. (B) Nonparametric dispersion test. (C) Zero-inflation test. (D) Q–Q plot used to assess deviations from the expected residual distribution, together with tests for uniformity (Kolmogorov–Smirnov), dispersion, and outliers. (E–F) Scaled residuals plotted against predicted values and against each model predictor. Simulation outliers—observations falling outside the range of simulated values—are highlighted as red stars, while red trend lines indicate statistically significant deviations from model expectations. Deviations were primarily associated with the predictor occurrence density, which likely reflects zero inflation arising from undersampled sites rather than true model misspecification. (G) Bivariate relationship between species richness and occurrence density.

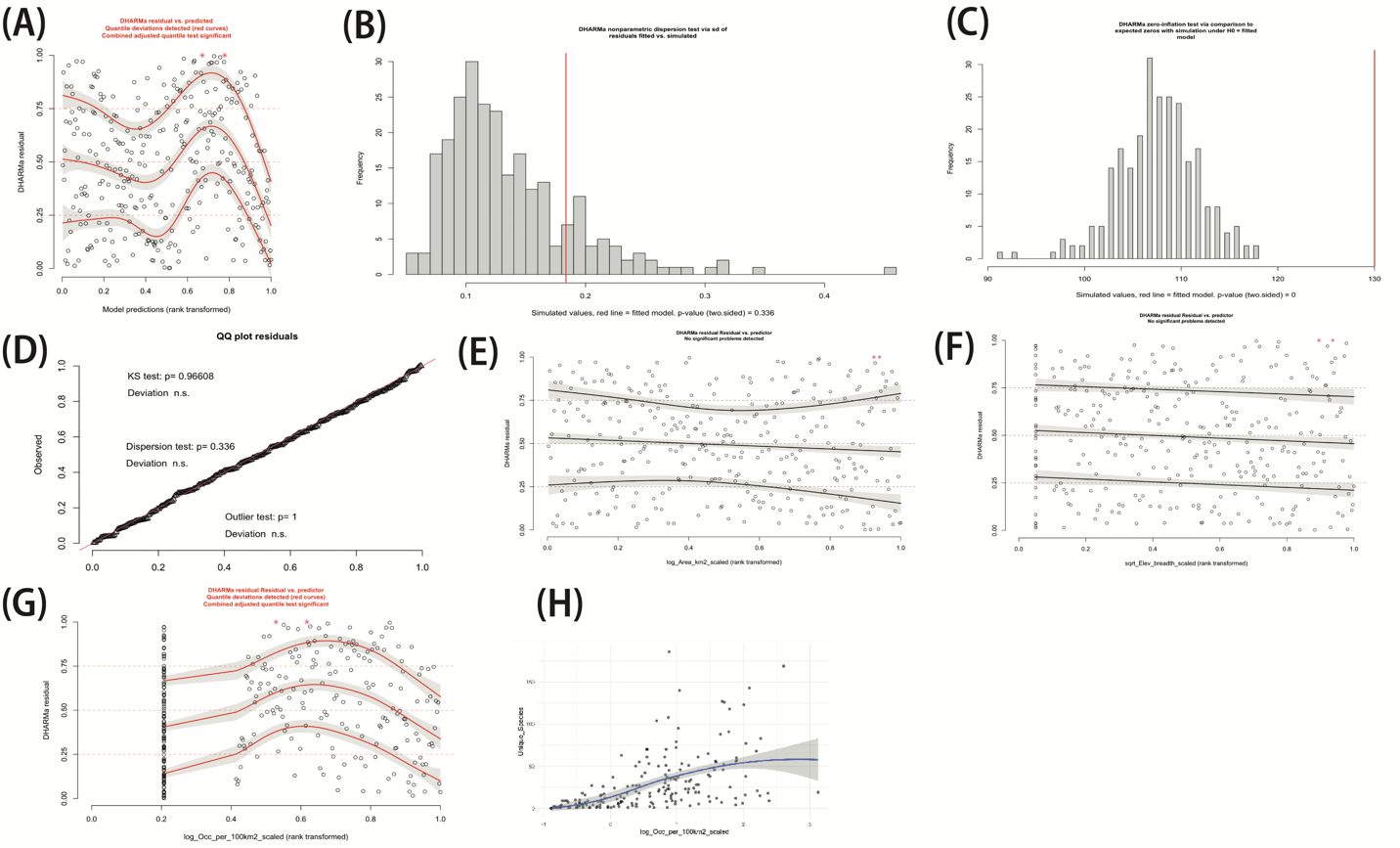

**S2 Fig. Model diagnostic results from DHARMa for the sampling effort model of Philippine conservation-relevant areas.** (A) Simulated residuals from the fitted model, showing detected quantile deviations. (B) Nonparametric dispersion test. (C) Zero-inflation test. (D) Q–Q plot used to assess deviations from the expected residual distribution, together with tests for uniformity (Kolmogorov–Smirnov), dispersion, and outliers. (E–F) Scaled residuals plotted against predicted values and against each model predictor. Simulation outliers—observations falling outside the range of simulated values—are highlighted as red stars, while red trend lines indicate statistically significant deviations from model expectations. (G–H) Bivariate relationship between species richness and each model predictor. Deviations were primarily associated with each predictor, likely reflecting increased variance at large PCAs where fewer observations exist.

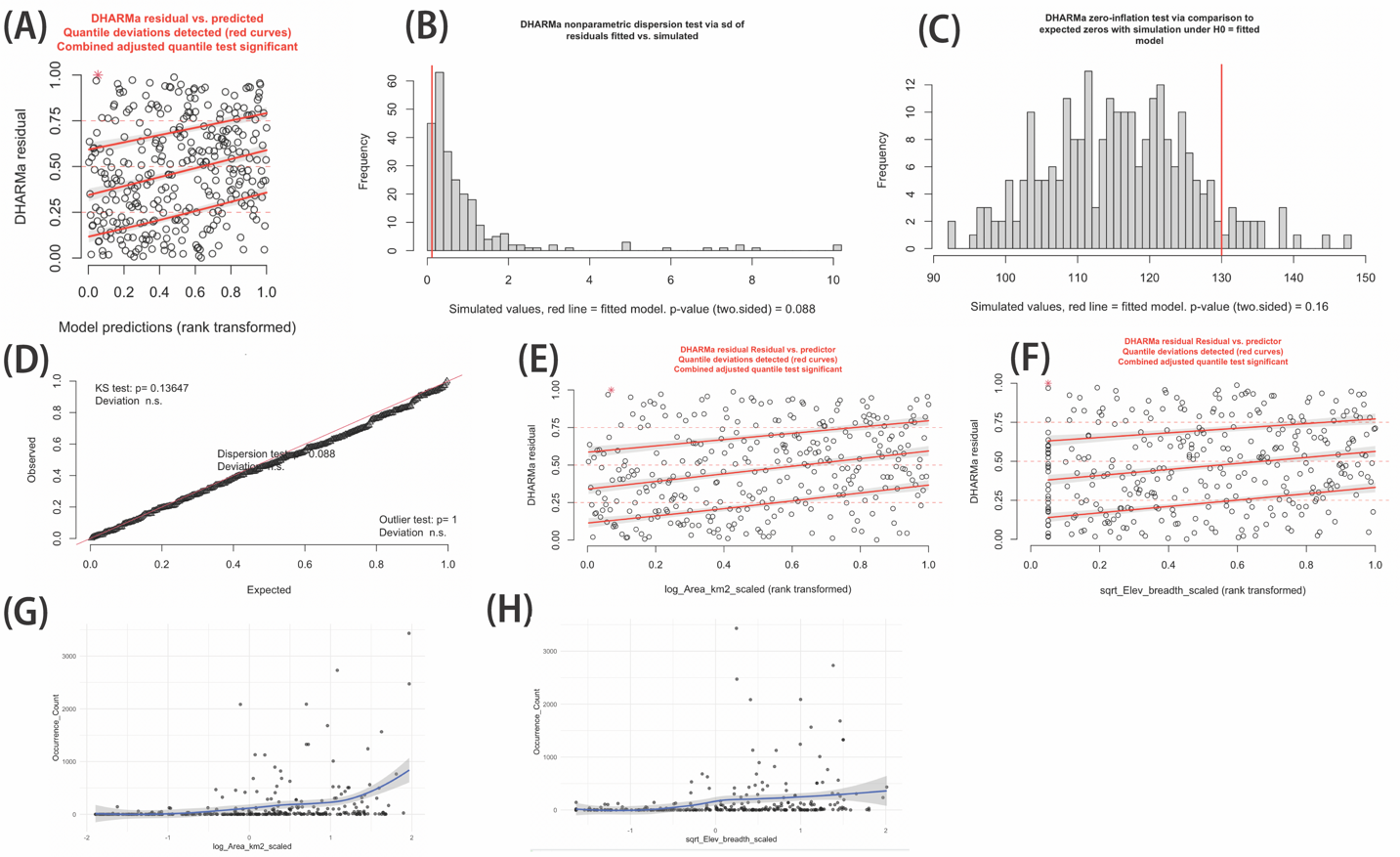

**S3 Fig.** **Map of the Philippines,** showing (A) overlap between forest cover and Philippine conservation-relevant areas (PCAs); (B) distribution of observed amphibian and squamate reptile diversity across all combined data sources outside PCA boundaries, highlighting that many well-sampled sites with higher observed species richness lie beyond established conservation-relevant areas. Administrative boundary and biodiversity area shapefiles were sourced from NAMRIA and the DENR-BMB via Geoportal Philippines (<https://www.geoportal.gov.ph/>).

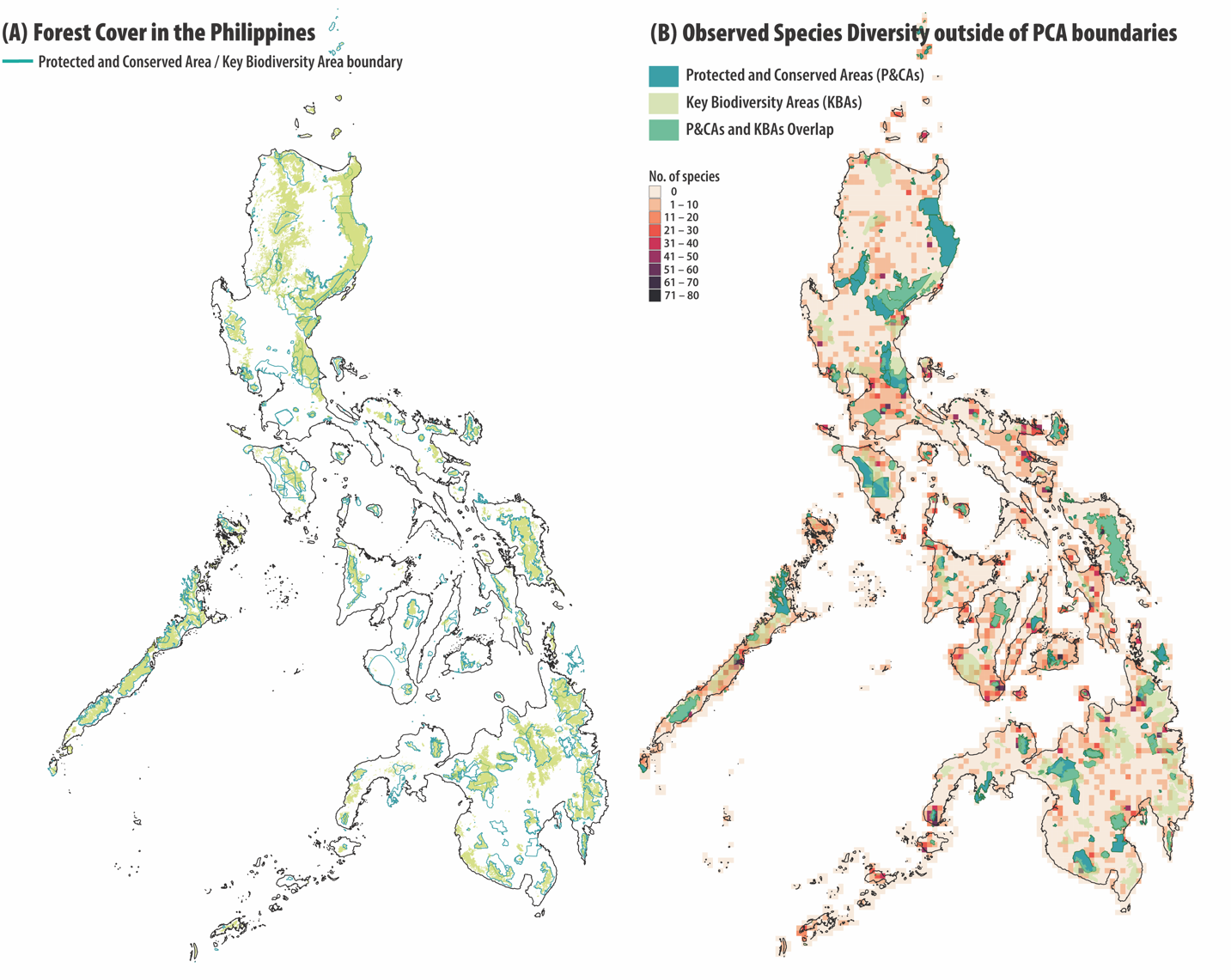
